# Identifying cell culturing parameters that improve endocytic uptake of the HIV-TAT cell penetrating peptide

**DOI:** 10.1101/2022.10.18.512764

**Authors:** Joshua Diaz, Jean-Philippe Pellois

## Abstract

Delivery tools, including cell-penetrating peptides (CPPs) are often inefficient due to a combination of poor endocytosis and endosomal escape. Herein, the impact of cell culturing techniques on the endocytic uptake of a typical CPP, the TAT peptide (derived from HIV1-TAT), was quantified. Parameters previously found to generally modulate endocytosis such as cell density, washing steps, and cell aging did not affect TAT endocytosis. In contrast, cell dissociation methods, media, temperature, serum starvation, and media composition all contributed to changes in uptake. The combination of these parameters in worst versus best-uptake protocols, led to changes in uptake of more than 13-fold and indicated that small variations in cell culturing techniques have a cumulative effect on CPP uptake. More specifically, modulating cell culture protocols does not result in an increased amount of peptide inside endosomes, rather the number of TMR-TAT containing endosomes increases. Taken together this study highlights how technical aspects of cell culture protocols can be used to improve experimental reproducibility, as well as parameters that can be potentially exploited to improve CPP accumulation in endosomes, and hence increase the possibility of endosomal escape and cytosolic access.

## Introduction

Many cellular delivery vectors utilize the endocytic pathway as a route to achieve cell penetration. In this context, cytosolic penetration can be thought of as a two-step process. First, delivery vectors and payloads are internalized by cells via various endocytic uptake mechanisms, a process that then leads to the distribution of this material in the lumen of various endosomal organelles [1]. Second, delivery vectors disrupt endosomal membranes to allow for their entry into the cytosolic space [2]. Taken together, efficient endosomal escape is thought to be the result of efficient entry and endosomal membrane disruption [3]. Because endosomal escape is the result of the two previously mentioned steps, improving either or both steps will likely lead to increased delivery efficiency. In principle, endosomal escape can be improved by increasing the intrinsic ability of delivery agents to permeabilize endosomal membranes. In addition, it can also be improved by increasing the amount of delivery agent that accumulates inside endosomes. As a matter of fact, a recent study has shown that the difference of cytosolic penetration between different CPPs can be attributed to differences in endosomal uptake [4]. Overall, it is reasonable to envision that increasing accumulation of delivery agents within endosomes, also increases the likelihood that it will achieve endosomal escape. Hence, a deeper understanding of the parameters that govern endocytic uptake of delivery vectors is needed to improve the efficiency of various delivery platforms.

One reagent that has been shown to rely on endosomal accumulation and subsequent endosomal escape is a CPP derived from the HIV-TAT protein, the TAT peptide (residues 49-57). TAT has been successfully used in many model organisms such as various tissue cultures lines as well as *in vivo*. TAT has also been tested in over 25 clinical trials [5]. Successfully delivered payloads include small molecules, peptides, proteins, nucleic acids, and nanoparticles [1, 3]. Despite its utility, the endosomal escape of TAT is generally poor, and the amount of peptide-payload entering the cytosol is estimated to represent less than 1% of what remains trapped inside endosomes. Thus, TAT is limited to payloads that do not require substantial amounts of molecules to exert their desired biological activity. Conversely, payloads that require higher concentrations of material to perform their biological activities may not be amenable for TAT-based delivery. Hence much effort has been expended on studying the mechanisms of TAT endosomal entry and release, which can aid in understanding how to improve delivery and ultimately to broaden the scope of molecules that can be delivered by TAT and TAT-like CPPs.

As the first step to enter the endocytic pathway, it is thought that TAT, which is arginine-rich and polycationic, interacts with cell surface anionic heparan sulfate proteoglycans [6-9]. The peptide is then thought to stimulate clustering of proteoglycans and its subsequent internalization by macropinocytosis [10]. This is typically demonstrated with the use of macropinocytosis inhibitors, such as amiloride, cytochalasin D, and rapamycin which can dramatically decrease TAT uptake (one caveat being that chemical inhibitors often lack specificity and may have pleiotropic effects), or by reduction in uptake from truncated proteoglycans that lack the ability to engage cytoplasmic partners [10-13]. Other endocytic uptake pathways are also potentially exploited by the peptide simultaneously [8, 14]. Nonetheless, because macropinocytosis leads to the formation of vesicles that are substantially larger than other endocytic processes (>250 nm macropinosomes vs <100 nm for other endocytic vesicles), it is likely that macropinocytosis contributes to the majority of CPP that accumulates inside cells [15, 16]. In fact, one study employed the use of luciferin-conjugated CPPs and endocytosis inhibitors to experimentally determine that macropinocytosis is the main route of entry for a variety of CPPs including TAT, with clathrin and cavaeolae/lipid raft dependent endocytosis playing minor roles in CPP uptake [17].

To determine the parameters that impact endocytic uptake and endosomal escape, many researchers have sought to establish structure-activity relationships that control TAT uptake into cells. They relate primarily to the arginine content of TAT, its concentration, and the presence of molecules that bind to TAT and prevent it from interacting with cell components [18-20]. In contrast, cellular and environmental factors that modulate TAT uptake are not fully understood. Given that the cell culture protocol used to deliver TAT could impact how TAT interacts with biological membranes and how biological functions associated with endocytic flux could be altered, we reasoned that it would be valuable to determine how different approaches to execute each step of the TAT delivery protocol can impact TAT endocytic uptake. We also aimed to identify parameters that would lead to consistent day-to-day endocytic uptake. These results would then potentially serve as the basis for protocols that lead to improved and reproducible CPP delivery outcomes.

## Results

### Experimental strategy and design

Many cell culture studies use adherent cells for cell biology and microscopy applications. Standard delivery protocols therefore include steps involving passaging, cell growth, washing, and incubation (figure 1). In order to assess the impact of these different processes on endocytic uptake, we labeled TAT with the fluorophore carboxytetramethylrhodamine (TMR) as a model peptide (TMR-TAT). We chose TAT to measure endocytosis because the endosomal escape of TAT is relatively poor. More efficient TAT-based delivery reagents typically display a mix of endosomal and cytosolically localization. Hence, TAT serves as an ideal reagent to decouple endocytosis from endosomal escape, as only a small portion (less than 1%) of the TAT-treated cells contain endocytosed TAT that has been released from endosomes.

**Figure 1:**
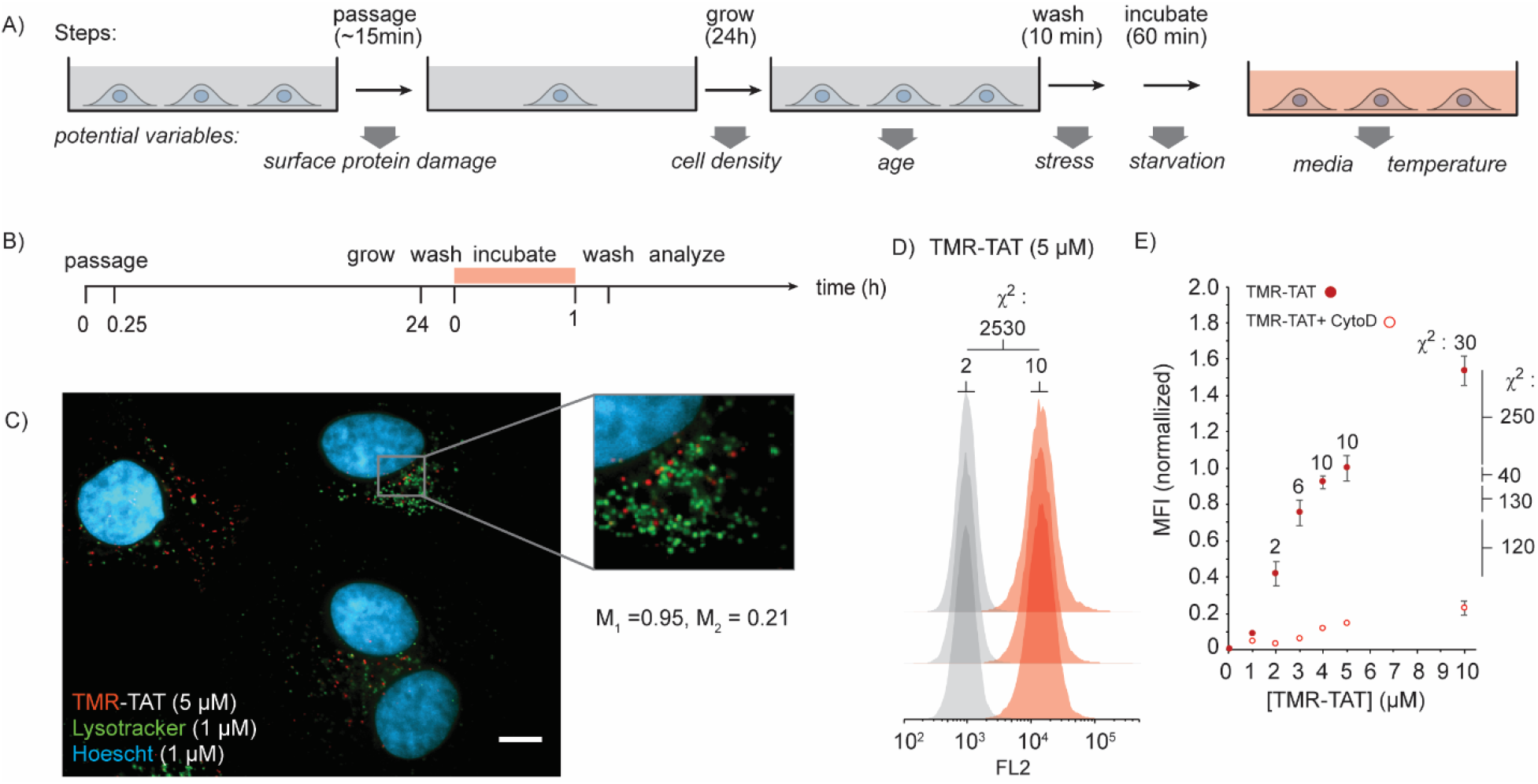
TMR-TAT as a probe to measure endocytic uptake. A) Scheme representing a standard protocol for a CPP-delivery experiment into adherent cells. The variables that may impact cellular uptake and that are tested in this report are highlighted. B) Scheme highlighting the timeframe chosen to perform TMR-TAT uptake assays. C) Representative fluorescence microscopy image of cells incubated with TMR-TAT (100x image, scale bar = 10 µm). The TMR-TAT signal is observed as red puncta that partially co-localizes with Lysotracker green (TMR-TAT is pseudocolored red, Lysotracker is pseudocolored green, and the nuclear stain Hoescht is pseudocolored cyan). A zoom-in image of the puncta is provided, along with the Manders coefficient M_1_ and M_2_ (M_1_ calculates the percentage of total signal from the red channel that overlaps with the green signal, and M_2_ calculates the percentage of green signal that is red). Colocalization is also observed by the color yellow which results from the overlay of green and red. D) Representative flow cytometry histograms of cells incubated with TMR-TAT (5 µM) for 1 h. Histograms are pseudocolored grey for untreated cells, and red for cells treated with CPP. FL2 is the fluorescence channel used to detect TMR. The histograms presented are for technical triplicates performed on the same day, using the same cell batch subjected to the same protocols. Chi squared (χ^2^) analysis is performed to compare the cell populations and establish statistical significance. E) Correlation between the amount of TMR-TAT (1-5 µM) incubated with cells and the corresponding median fluorescence intensity (MFI) of the cell populations detected by flow cytometry. The data presented are the average and corresponding standard deviations obtained from biological triplicates (same protocol, but different cell batches on different days), normalized to the TMR-TAT 5 μM condition. The Chi squared numbers reported above each data point compare the results obtained from biological triplicate (the highest χ^2^ obtained being reported). The Chi squared numbers reported on the right side compare the samples that used different CPP concentrations (the lowest χ^2^ obtained being reported). The flow cytometry results obtained in the presence of CytoD (20 µM), an inhibitor of actin polymerization and of macropinocytosis are also provided.

The cell line used is HeLa because it is one of the most commonly utilized cell lines and its use has been cited in over 60,000 publications [21]. Incubation of TMR-TAT with HeLa for 1 hour leads to the formation of fluorescent puncta. These puncta colocalize with lysotracker green, indicating that TMR-TAT is present within the lumen of endosomes (Figure 1C). The total fluorescence intensity of cells was measured by flow cytometry. Cells incubated with TMR-TAT typically display a unimodal bell shape distribution (Figure 1D). Experiments were performed with technical triplicates (experiments performed on the same day with 3 different tissue culture dishes) and flow cytometry data was analyzed with the FlowJo software. In particular, the Chi squared T(x) score (χ^2^ in the figures) was used to estimate the probability that two populations are different from one another [22-24]. The Chi squared T(x) score between untreated cells was 2 or less, and 10 or less for TMR-TAT incubation triplicates (Figure 1D). Titration experiments show a positive correlation between the concentration of TMR-TAT during incubation (from 1 to 10 μM) and the median intensity of cells detected by flow cytometry. The Chi squared T(x) scores between triplicates of the same conditions were 30 or less. In contrast, the Chi squared T(x) scores between populations representing different TMR-TAT concentrations were between 40 and 250. Overall, for subsequent experiments, we consider populations with a Chi squared T(x) score of 40 or less as not statistically different while populations with a Chi squared T(x) score of 40 or more are considered statistically different (the confidence of this assessment increasing with the value of the Chi squared T(x) score). Please note that biological triplicates (the same experiment performed on 3 consecutive days), yielded Chi squared T(x) score that exceed 40. This variability is in part the issue surveyed in this report. Additionally, for all subsequent experiments, we use the standard protocol (SP), changing only one variable at a time. We use results obtained from technical triplicates performed on the same day to compare the impact of individual parameters. In contrast, experiments performed on different days are not compared to one another. Microscopy experiments were conducted in parallel of all flow cytometry analyses to confirm that the bulk of the fluorescent signal detected by flow cytometry corresponds to fluorescent endosomes. Furthermore, the influence of cell aging on the variability of the biological replicates was tested. This was done using various stocks of HeLa cells that were cultured for various timescales (2 days, 10 days, 25 days) and stored in a liquid nitrogen dewar. On the same day cell stocks were revived, passaged in parallel and then tested for uptake on the same day. It was found that cell aging within this 25-day time frame had no effect on endocytosis (figure S1). All assays conducted did not use HeLa cells cultured for 25 days past the original stock.

**Figure S1:**
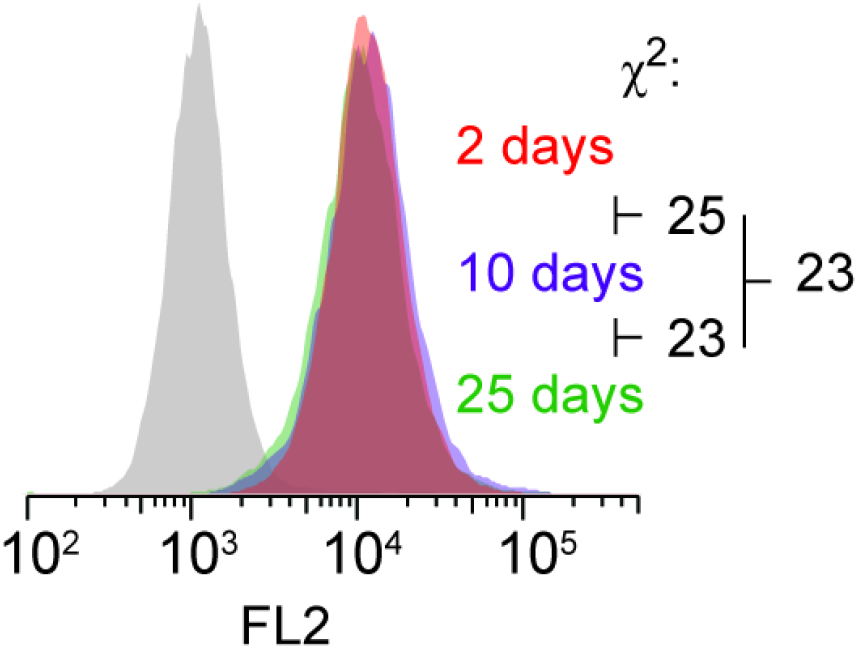
Age of HeLa cells does not impact TMR-TAT uptake. HeLa cells were grown in a 37°C incubator and passaged for up to 25 days. At the 8^th^ day and 23^rd^ day mark, cells were stored in a liquid nitrogen dewar. All cell stocks (previously cultured 0 days, 8 days, and 23 days) were then revived at the same time and passaged twice. Cells (passaged 2, 10, 25 days respectively) were then treated with TMR-TAT (5 µM) for 1 hour and subjected to flow cytometry analysis.

### Impact of passaging on endocytic uptake of TMR-TAT

A standard step involved in maintaining cells and preparing them for a new experiment is to transfer them to a new dish. For adherent cells, this step involves detaching cells from the current dish in which they are grown. This can be performed by addition of a low amount of the protease trypsin (0.05%-0.5%), which cleaves cell surface proteins [25, 26]. Alternatively, proteins involved in cell-to-cell adhesion and extracellular matrix attachments can be disrupted. This can be performed because they require metal ions such as Ca^2+^, Mn^2+^, or Mg ^2+^ for proper function. Hence, enzyme-free cell dissociation buffers (CDB) that contain chelators such as EDTA can be used to release cells from a surface [27-29]. Notably, by modifying cell surface proteins to different extents, both methods can potentially impact endocytic uptake. To test this variable, the incubation protocol described in Figure 1 was applied to cells, with the exception that cells were either dissociated from a growth dish with 0.5% trypsin or with CDB. Cells were then added to new dishes and incubated for 4 hours or 24 hours (4 hours being the shortest time required to for a majority of cells to recover cell adhesion). Cells were then incubated with TMR-TAT and endocytic uptake was measured by flow cytometry. In parallel experiments, cell-surface proteins were labeled with cell-impermeable NHS-fluorescein. Cell lysates were then analyzed by SDS-PAGE and fluorescein labeled proteins were detected by fluorescence scanner. As expected, a 5 minute trypsin treatment leads the proteolytic degradation of labeled proteins, as indicated by loss of high molecular weight bands with a fluorescein signal on SDS-PAGE (figure 2). In contrast, the fluorescent band pattern was not impacted by treatment with CDB for 5 minute. High molecular weight bands are recovered 4 hours post-treatment for the trypsin condition, albeit with a pattern noticeably distinct from either the untreated or CDB conditions. The flow analysis of cells treated with CDB and incubated with TMR-TAT at either the 4 hours or 24 hours post-treatment time points, show a high reproducibility. In particular, the median fluorescence intensities (MFI) between replicates are comparable between the 4 hour and 24 hour time points (as established by Chi squared T(x) score of 16). Overall, these results are consistent with the notion that CDB does have a deleterious effect on the cell surface and as a result, endocytic uptake is relatively unaffected by this treatment. In contrast, TMR-TAT endocytic uptake was highly variable in trypsinized cells. This is observable at both the 4h and 24h time points Chi squared T(x) score of 100).

**Figure 2:**
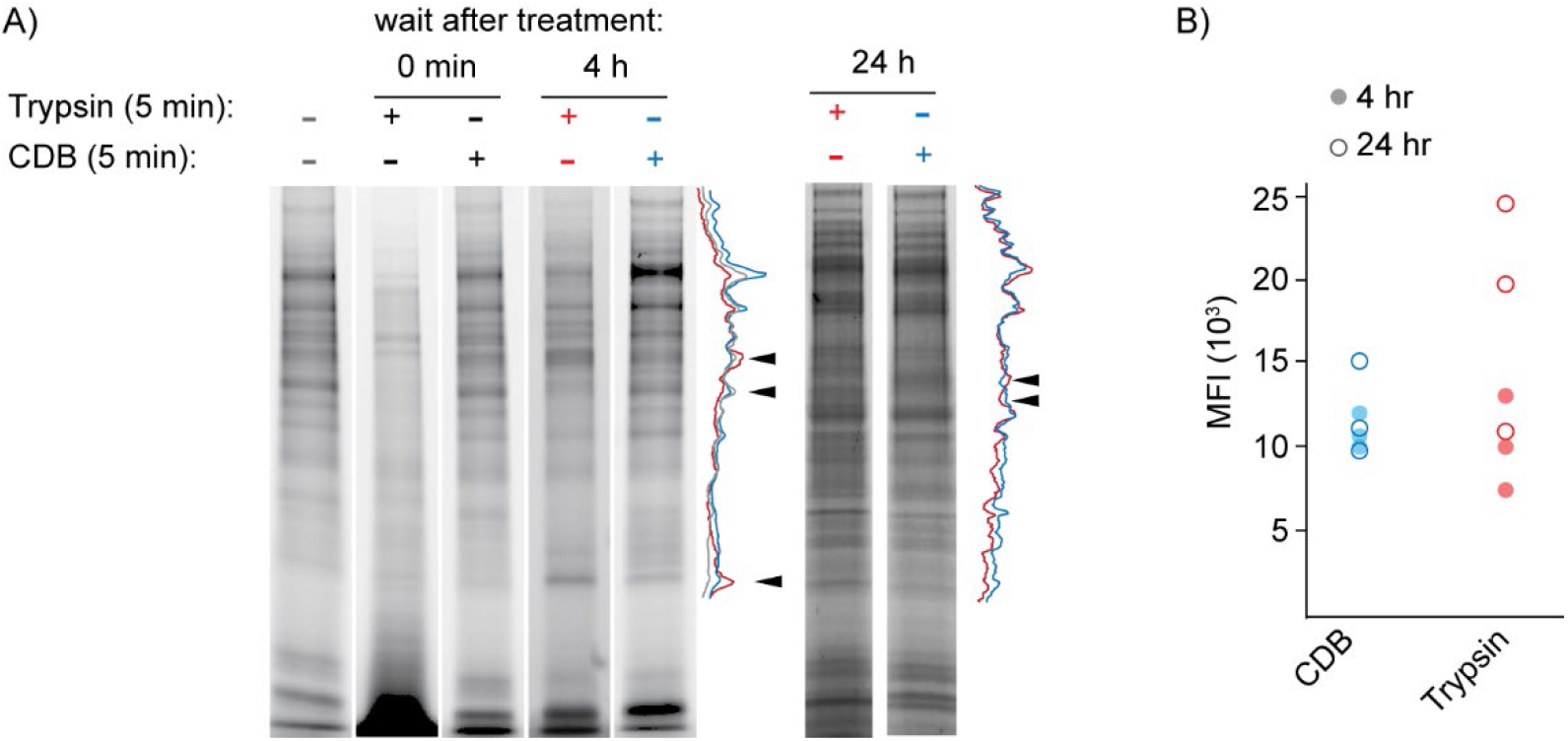
Cell passaging impacts TMR-TAT uptake. A) SDS-PAGE analysis of cell surface proteins labeled with NHS-Alexa Fluor™ 488. Cells were labeled and analyzed immediately after treatment with 0.5% trypsin or cell dissociation buffer (CDB), or after 4 or 24 h of recovery time. Line profile intensities of samples obtained after a 4 or 24 h recovery (pseudocolored grey for untreated cells, pseudocolored red for cells treated with trypsin, and pseudocolored blue for cells treated with CDB) are shown. Black arrows highlight band intensities that differ between samples. B) Flow cytometry MFI obtained from cells incubated with TMR-TAT (5 μM, 1h) 4 or 24 h after treatment with trypsin or CDB. Each data point is obtained from cells cultured and passaged on different days.

### Cell density

The density of cells adhering to a surface, or whether cells make contact with one another, has been reported to impact entry of different viruses and endocytic uptake of fluorescent transferrin [29]. In order to test how this impacts the uptake of TMR-TAT, cells were seeded at different dilutions during passaging and allowed to grow overnight. To quantify cell density, cells were stained with Hoechst and imaged by fluorescence microscopy (over the whole area of the dish to ensure homogeneity of distribution). Confluent cells were used to establish a maximal density of blue nuclei. This was then used to determine a ratio of surface density (Figure S2). Cells were also stained with di-4-ANEPPDHQ, a lipophilic probe, to label their membranes and to confirm that cells contact each other at high density [30]. Cells at various densities were then incubated with TMR-TAT using the SP and analyzed by flow cytometry. The populations of TMR-TAT positive cells were indistinguishable in all conditions tested, as indicated by Chi squared T(x) scores of zero (Figure S2).

**Fig S2:**
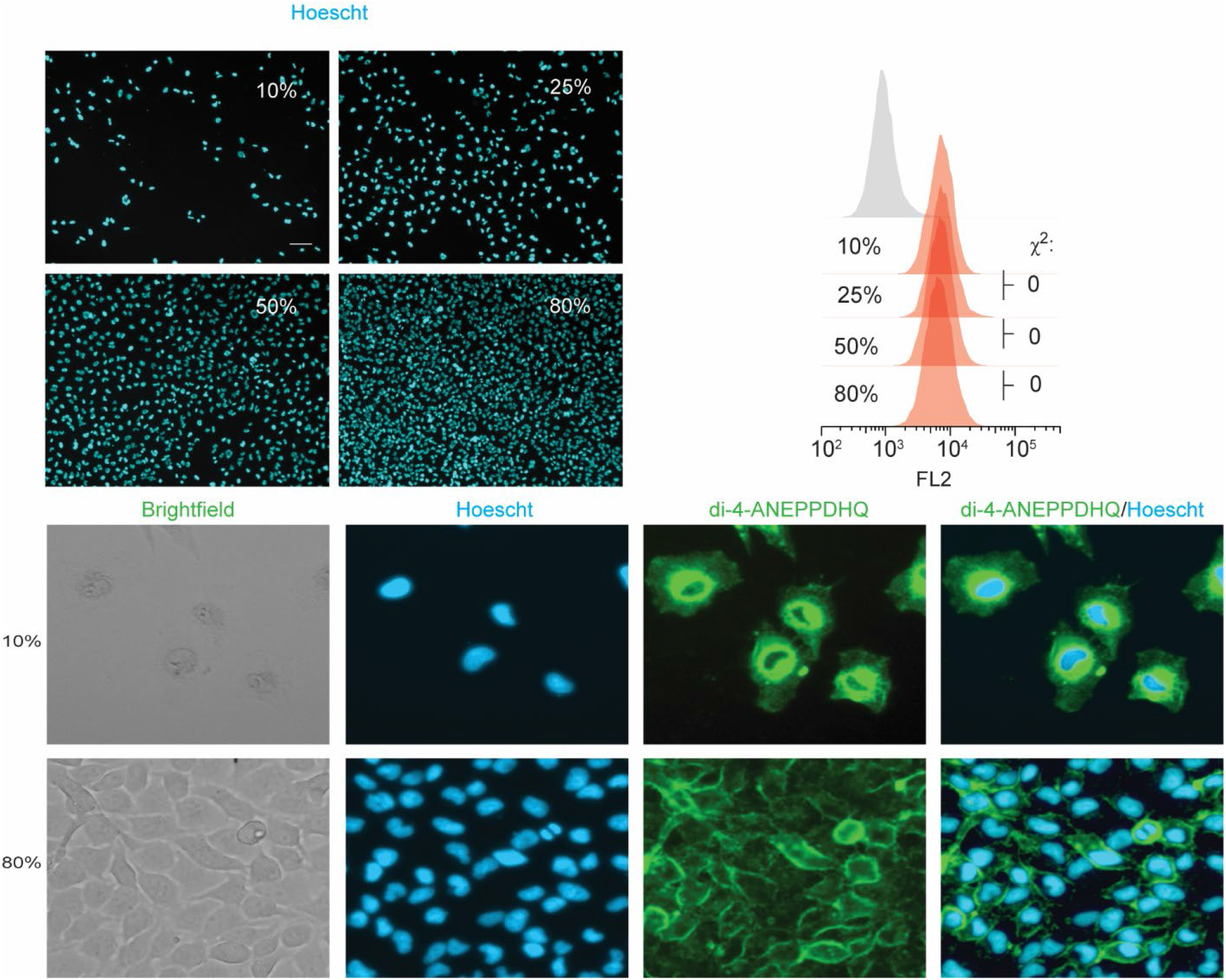
Cell density does not impact TMR-TAT uptake. A) Cell density measured by cell nuclei staining (scale bar 150 uM, 10x image). Cells were plated at various confluences by cell counting on a hemacytometer. B) Quantification of TMR-TAT uptake at different cell densities. Cells were treated with TMR-TAT (5 µM) for one hour. C) Cell surface staining via lipophilic di-4-ANEPPDHQ. 50 µM scale bar 40x image. We attribute the differences in di-4-ANEPPDHQ staining between the 10% and 80% images differences in morphology observed in brightfield images.

### Culturing cells

In a typical tissue culture experiment, cells are cultured in a media adequate for growth and proliferation. Cells can then be incubated in a media better adapted to a delivery protocol and to a desired application. In the case study described herein, this involves removing phenol red-DMEM supplemented with 10% FBS, which is a medium using a sodium bicarbonate buffering system adapted to growth in 5% CO2 incubators, and replacing it with L-15, a medium buffered by phosphates and compatible with ambient air (and hence better suited for microscopy experiments). The L-15 used is usually serum free, as FBS, and albumin in particular, are thought to interfere with CPPs [31]. This is likely because albumin binds to cationic CPPs and thereby competes with their interactions with cell components. In fact, several cationic antimicrobial peptides have been demonstrated to bind albumin [32, 33]. Despite being relatively trivial, media exchange can be stressful to cells. In particular, flow and shear stress can impact cell physiology, both in vivo and in vitro [34-37]. Flow cytometry assays however indicated that different washing protocols did not impact the endocytic uptake of TMR-TAT (despite affecting cell adhesion and viability) (Figure S3). Likewise, the number of repeated washing steps (from 1 to 3) did not have an effect on flow cytometry results (with the exception of 0 wash, where uptake is inhibited possibly because of unwashed BSA) (Figure S4). Notably, the uptake of TMR-TAT was influenced by the media used during TMR-TAT 1 hour incubation. In particular, endocytosis of TMR-TAT was higher in DMEM (serum free DMEM) than in L-15 by an approximate factor of 1.4 (Figure 3A). Beyond differences in buffering component composition, DMEM and L-15 differ in oxidizable carbon source, DMEM being formulated with low or high glucose (5 or 25 mM), and L15 being formulated with galactose (5 mM) and sodium pyruvate (5 mM). These differences can contribute to metabolic reprogramming (i.e. production of less lactate, lower proliferation rate in galactose vs glucose media) and potentially impact the regulation of endocytosis.[38, 39] In order to assess the effect of carbon sources on TMR-TAT uptake, we first tested whether the medium used for cell growth has an impact on endocytosis. In this experiment, cells were grown in either high or low glucose DMEM (HG-DMEM, 25 mM; LG-DMEM 5 mM) for 24 hours. Cells were then incubated with TMR-TAT in either L-15 or HG-DMEM for 1 hour. A difference in uptake was not detectable in the L-15 condition, and small in the DMEM samples (Chi square T(x) score of 55) (Figure 3B). Moreover, the growth media did not change the difference observed between L-15 and HG-DMEM peptide incubation media. To test whether glucose is one of the media components that influence endocytic uptake, cells were incubated with peptide in DMEM with or without glucose (25 or 0 mM) and in L15 with or without glucose (25 or 0 mM glucose, 5 mM galactose). Removing glucose from DMEM during the 1 h incubation caused a reduction in TMR-TAT uptake. Conversely, adding glucose to L15 increased the uptake of TMR-TAT (Figure 3C). Overall, these results indicate that glucose stimulates the endocytic uptake of TMR-TAT. However, because differences in uptake remain between L15+glucose and DMEM+glucose, other media components may also play a role.

**Fig S3:**
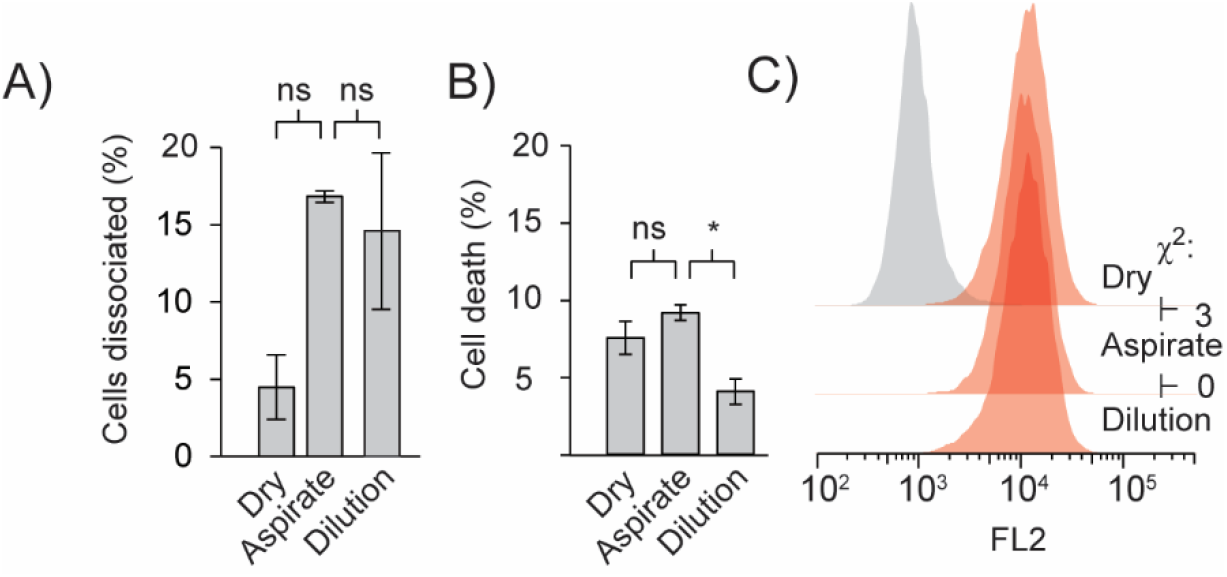
Potential cell stress resulting from washing techniques do not impact TMR-TAT uptake. A) Quantification of dissociated cells. Cells were stained with Hoescht (1 µM) and the entire well was imaged using an automated EVOS-M7000. The resulting images were then stitched using the EVOS7000 stock software and stained nuclei were subsequently counted using Celleste Software. The cells were then subjected to washing. After washing the cells using different techniques, they were imaged again and counted as previously described. Both cell counts allowed for a determination of cell dissociated (%). This experiment was repeated in triplicate and the t-test was performed to analyze significance. B) Quantification of cell death. Cells were washed using different techniques. Cell death was established by determining the total cell population as in (A) and subsequent incubation with DRAQ7 (1 µM). C) Quantification of uptake of TMR-TAT post different washing procesdures. Cell were washed using different washing techniques. Subsequently, cells were treated with TMR-TAT (5 µM) and analyzed by flow cytometry.

**Fig S4:**
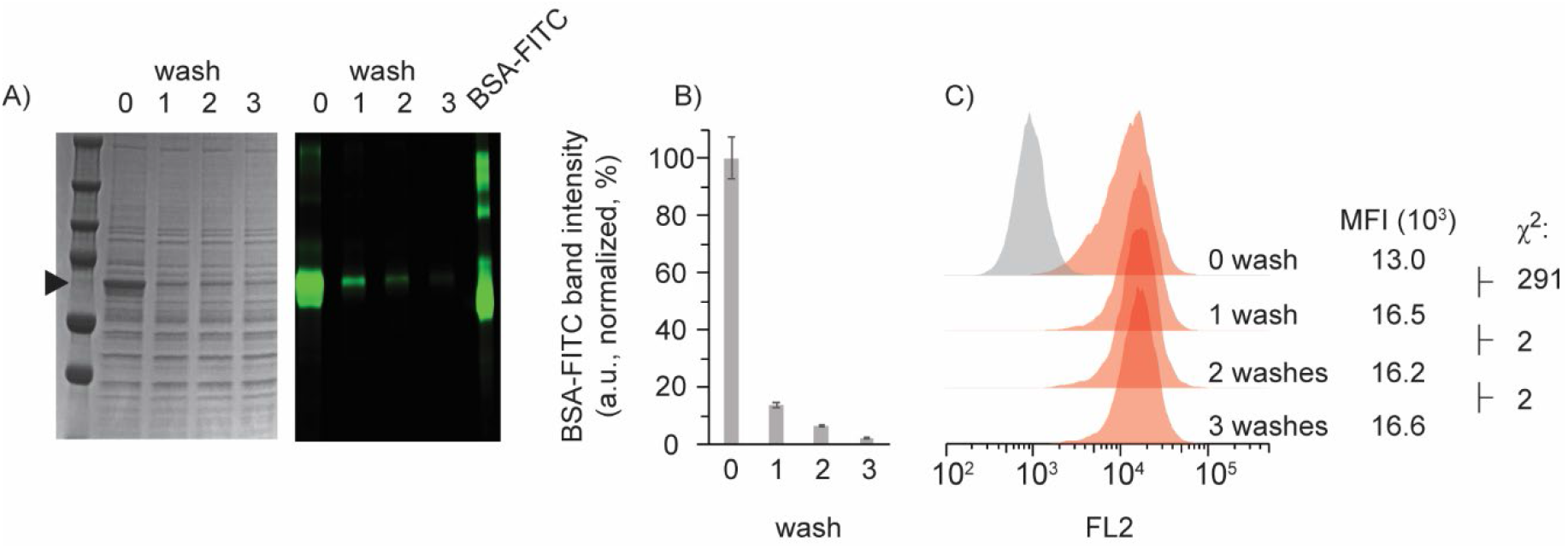
Serum supplementation in DMEM growth media hinders TMR-TAT uptake. A) SDS-PAGE analysis of removal of BSA-FITC from growth media. BSA-FITC (3.25 µM) was incubated for 24 hours in DMEM supplemented with 10% FBS. Cells were then washed 0, 1, 2, 3 times. Cells were then lysed and lysates were loaded onto SDS-PAGE. Purified BSA-FITC was loaded onto SDS-PAGE for comparison and BSA-FITC was then scanned for fluorescence. B) Quantification of relative amounts of BSA remaining. Each band representing BSA-FITC was analyzed by densitometry and subsequently normalized to a purified BSA-FITC that had similar concentration to the 0 wash. This experiment was performed in duplicate. C) Cells were washed either 0, 1, 2, or 3 times. Cells were then treated with TMR-TAT (5 µM) for 1 hour and analyzed via flow cytometry.

**Figure 3:**
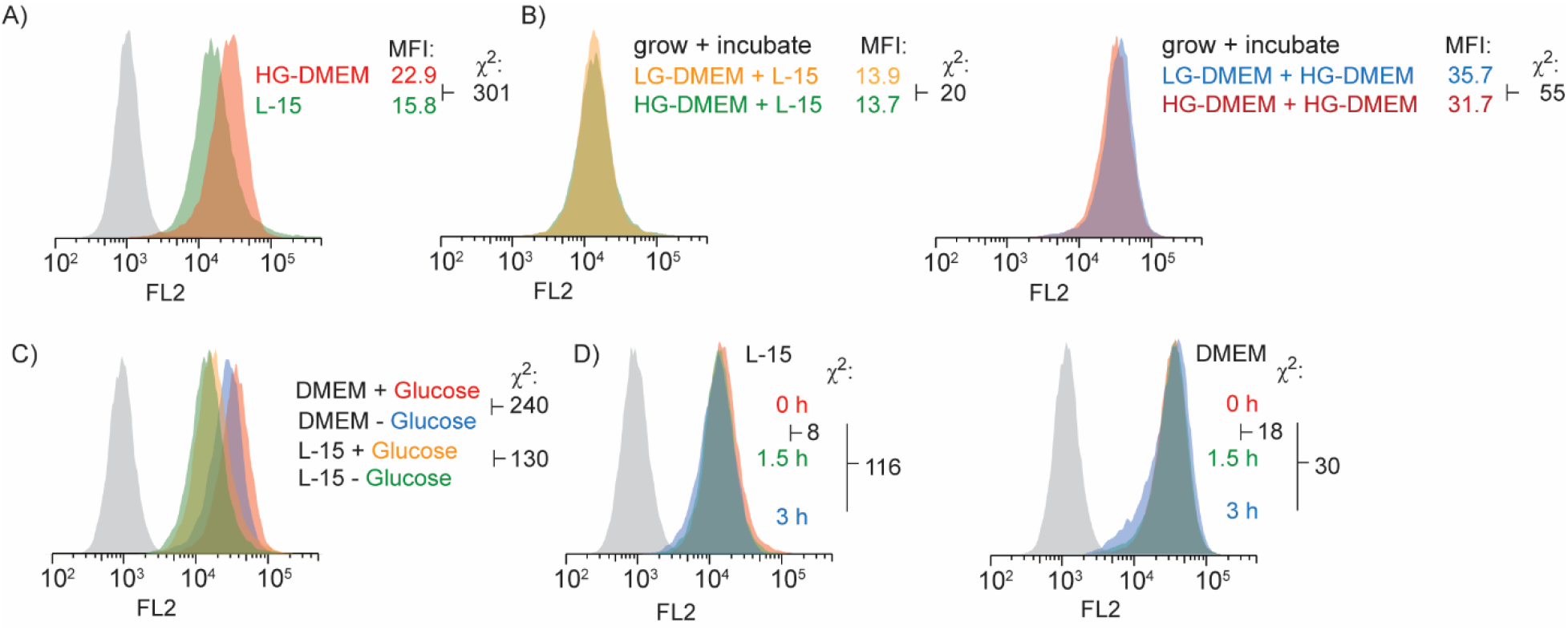
Impact of growth and incubation media on TMR-TAT uptake. A) Representative flow cytometry histograms of cells incubated with TMR-TAT (5 μM, 1h) in either HG-DMEM or in L-15. B) Representative flow cytometry histograms of cells grown for 24 h in either LG-DMEM (5 mM glucose) or HG-DMEM (25 mM glucose), and incubated with TMR-TAT (5 μM, 1h) in either HG-DMEM or in L-15. C) Representative flow cytometry histograms of cells incubated with TMR-TAT (5 μM, 1h) in DMEM or in L-15 with or without glucose added (the glucose concentration is 0 or 25 mM; DMEM with 25 mM glucose is HG-DMEM). D) Representative flow cytometry histograms of cells incubated in serum free L-15 or serum free DMEM for 0, 1.5, or 3h, and then incubated with TMR-TAT (5 μM, 1h) in L-15 or DMEM. All experiments were performed in technical triplicate and the Chi-square was determined to be 10 or less for all replicates (not shown). The media conditions and the corresponding histograms are pseudocolored with similar colors. The grey histograms are from cells not treated with TMR-TAT.

### Serum starvation

The change between serum supplemented media to a serum-free media is a common practice in molecular biology. The rationale for this procedure can vary depending on the goal of the cell-based studies, such as cell cycle synchronization or induction of cell stress responses [40-43]. In the context of CPPs, as mentioned above, FBS components such as albumin can compete with cell surface molecules for CPP binding. In our experience, albumin is inhibitory to cell penetration for TAT analogs [31, 44]. Additionally, serum starvation can elicit changes in the endocytic pathway and potentially impact CPP uptake indirectly [45]. To our knowledge, the effect of serum starvation on CPP endocytic uptake has however not been evaluated. To determine the effect of serum starvation on TMR-TAT uptake, cells were grown in 10% FBS HG-DMEM for 24 hours and switched to SF-L-15 or SF-HG-DMEM for 0, 1.5, or 3 hours. Cells were then treated with TMR-TAT in either of the two media for 1 hour. There was a notable decrease in uptake in cells starved in L-15. Conversely, cells starved in DMEM has little to no effect.

### Effect of Temperature on endocytic uptake rate

Temperature is an important variable in numerous cellular processes, including endocytosis. Particularly, macropinocytosis involves actin polymerization, which is temperature dependent [46]. While it is easy to maintain a temperature of 37°C when cells are in an incubator, it is less trivial to achieve during manipulations that take place in biosafety cabinets at room temperature. In principle, steps such as cell washes, media exchange and addition of CPP mixtures may be done with solutions which temperature fluctuate between 37°C and 25°C. In order to assess the effect temperature variation on CPP uptake, cells were incubated with TMR-TAT at either 37°C and 25°C for 5, 15, 30, 45 and 60 minutes (figure 4). The uptake of the peptide was consistently lower at 25°C than at 37°C. The median fluorescence intensities between 5 and 60 minutes fit a simple linear regression (if the time 0 data point is ignored), and the apparent rate of uptake at 37°C is approximately twice that observed at 25°C. At the 60 min incubation time point (condition used for all other experiments), the uptake of TMR-TAT is 1.9-fold less at 25°C than at 37°C.

**Figure 4:**
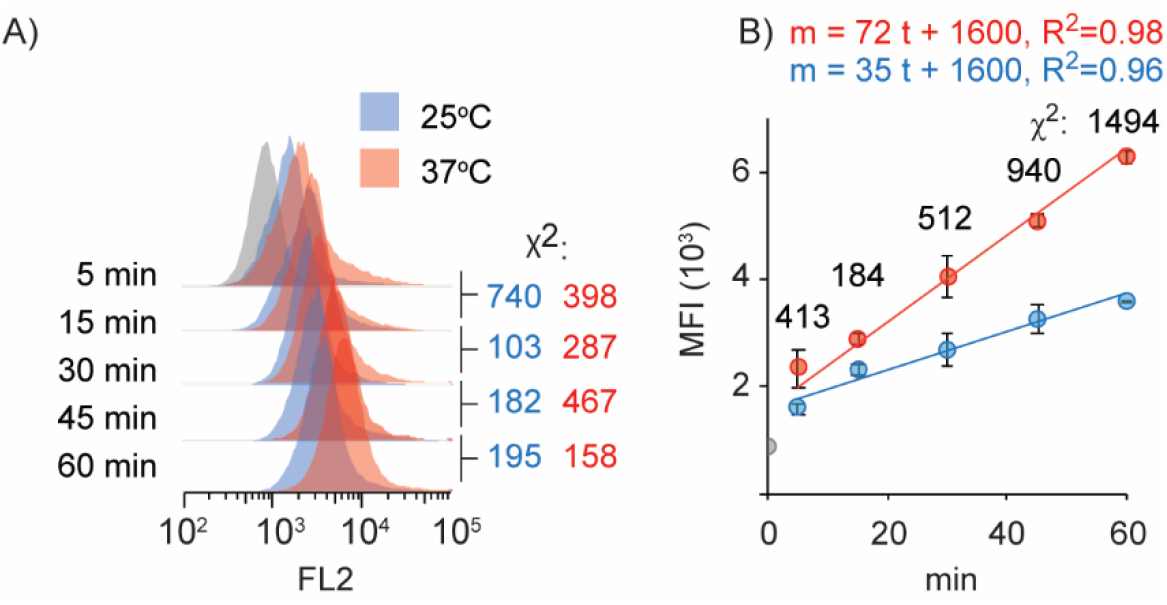
Impact of temperature on TMR-TAT uptake. A) Representative flow cytometry histograms of cells incubated with TMR-TAT (5 μM) in L-15 for 5-60 min, at either 25°C (pseudocolored blue) or 37°C (pseudocolored red). The chi-square values obtained between populations incubated for successive time points are provided. B) Graphs representing the change in FL2 MFI, as determined by flow cytometry, as a function of incubation time with TMR-TAT (5 μM). The red and blue graphs are obtained from cells incubated at 37°C and 25°C, respectively. Each data point is the average of technical triplicates, with corresponding standard deviations. The Chi-square values shown are obtained from comparing the populations analyzed at 37 and 25°C for each time point. Linear fit (which exclude time 0) and their corresponding equations and R^2^ are shown.

### Cumulative impact of protocol variables

Several individual parameters modulate TMR-TAT uptake, some by a relatively small amount (growth media, serum starvation), others having a more substantial effect (temperature, incubation media formulation). One can envision that different investigators may choose different combinations of these parameters for their protocols. We were therefore interested in testing the range of endocytic uptake that these different combinations could generate. Rather than testing all possible combinations, we choose to test three protocols. The first protocol combines the individual parameters that caused a reduction in uptake in the experiments described above (growth in HG-DMEM, 3h SF-L15 starvation, incubation in L15 at 25°C). This protocol is referred to as “low” uptake protocol (LUP). A medium uptake protocol (MUP) was designed to include the LUP factors, with the exception of including incubation at 37°C as opposed to 25°C. Conversely, a “high” uptake protocol (HUP) was designed to combined the optimal conditions identified in prior experiments, i.e. growth in LG DMEM, no serum starvation, incubation in HG-DMEM at 37°C. The difference between CDB and trypsin passaging could also in principle contribute to increasing or reducing uptake. However, given that trypsin gave non-reproducible and aleatory results, we did not use this parameter in this experiment. Instead, all samples were treated with CDB. Cells, prepared for the three uptake protocols, were treated with TMR-TAT (5 μM) for 1h. Flow cytometry analysis of the LUP cells population yielded a median fluorescence intensity (MFI) of 2.6 × 10^3^, the MUP cells yielded a MFI of 17.0 × 10^3^, and the HUP cells yielded a MFI of 37.5 × 10^3^ (Figure 5B). Using a median of 0.8 × 10^3^ for untreated cells, the HUP contributes to 13-fold and 2.8-fold increases in uptake when compared to the LUP and MUP, respectively. Moreover, the HUP yielded more uptake of TMR-TAT at 1 μM than the LUP at 5 μM, indicating that the change in incubation protocol is equivalent to a reduction in peptide concentration of 5-fold or more. Fluorescence microscopy imaging confirmed that with both protocols the peptide localizes in puncta consistent with endosomes. Interestingly, the average pixel intensity for these HUP and MUP puncta are not statistically different (Figure 5C), indicating that the amount of TMR-TAT molecules inside endosomes is equivalent for these conditions. In contrast, the number endosomes that are present per cell (quantified indirectly by counting the total number of pixels from TMR-TAT puncta; see Material and Methods for details) is on average 2-fold greater for HUP than MUP. Overall, we conclude that the HUP condition leads to approximately twice the amount of TMR-TAT uptake as the MUP condition. However, this is not because endosomes have accumulated twice the concentration of TMR-TAT in their lumen. Instead, our data suggest that this is because HUP cells have twice the number of TMR-TAT containing endosomes.

**Figure 5:**
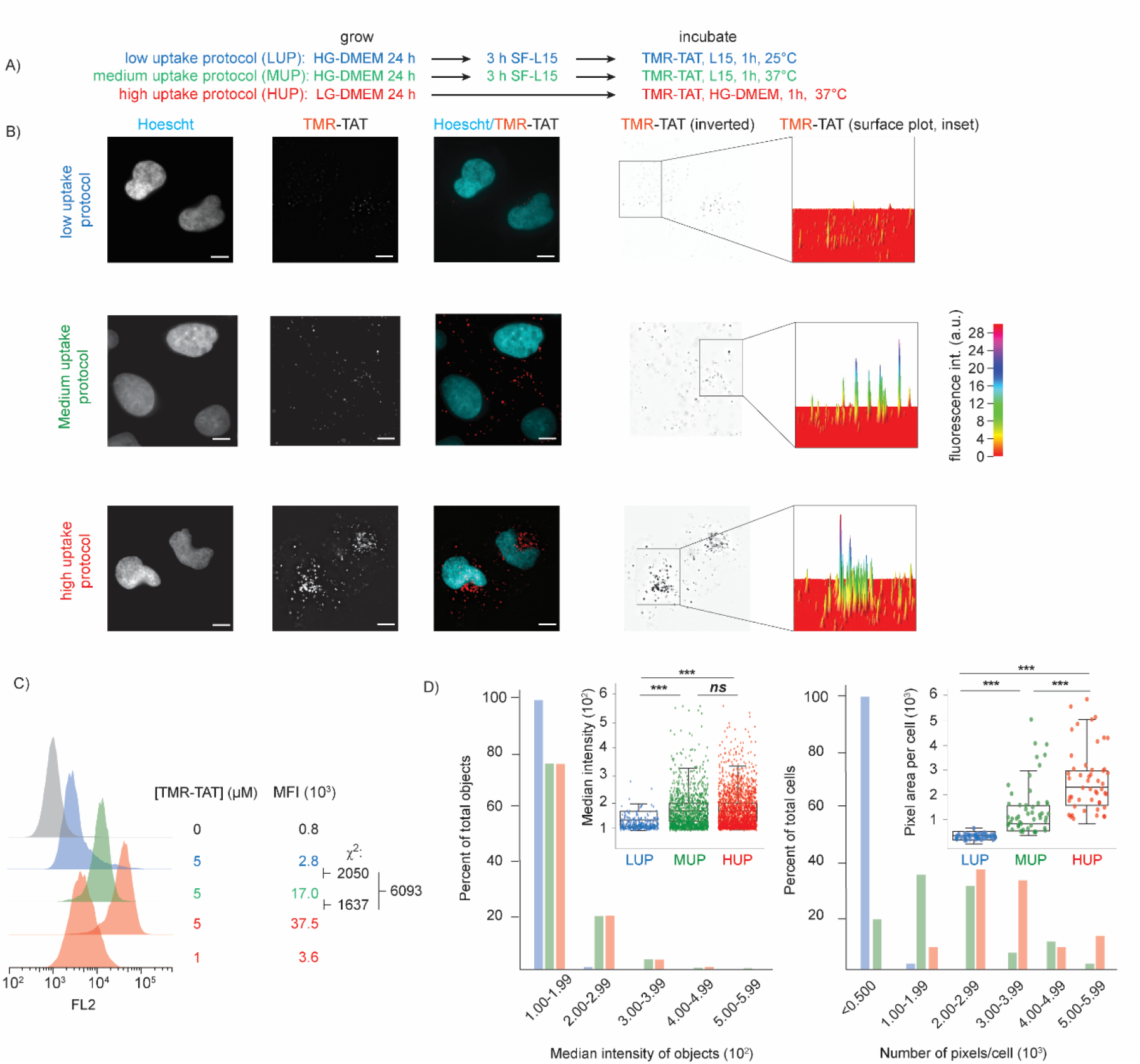
The effects of growth media, serum starvation, incubation media and temperature on TMR-TAT uptake are additive. A) Design of low (blue), medium (green), and high uptake (red) protocols based on previous results. B) Representative fluorescence microscopy images of cells (100x image, scale bar = 2.5 µm) treated with TMR-TAT using low, medium, and high uptake protocols. Imaging and image processing were performed using identical conditions. Images are monochromes for TMR-TAT, or pseudocolored cyan for Hoescht and red for TMR-TAT for overlay images. The 3D surface intensity plots for insets of the TMR-TAT images are provided. The signal intensities are displayed as a color gradient. C) Representative flow cytometry histograms of cells treated with TMR-TAT using low (blue), medium (green), and high (red) uptake protocols. The MFI reported are the average of technical triplicates. D) Quantification of TMR-TAT puncta, specifically pixel intensity (left) and pixel area per cell (right) for the low uptake (blue data points), medium uptake (green data points) and the high uptake (red data points) protocols. Fluorescence microscopy images were deconvoluted in Slidebook 6. In Celleste image analysis software, individual cells were selected (at least 50 cells counted for analysis). Puncta were then analyzed for both pixel intensity (each individual puncta) and pixel area (per cell). Data points for each analysis are displayed in a scatter plot (inset). These data points were then binned and are displayed in histograms.

## Discussion

Herein, we were interested in understanding how the endocytosis of a model CPP (i.e. a delivery vector likely representative of other delivery platforms that primarily utilize macropinocytosis for cell entry) is impacted by various steps associated with culturing protocols. More specifically, we also focused on parameters that are not necessarily biologically relevant (e.g. temperature sensitivity) but which are relevant from a technical point of view instead. Our rationale is that this foundational knowledge about cell handling is a prerequisite to potentially make *in vitro* tissue cultures better models for *in vivo* delivery studies (despite obvious gaps in complexity). Because *in vitro* cultures are also used to produce biologics or therapeutic in cells, we also reasoned that understanding technical parameters may also be of practical value in this context. Moreover, tissue cultures are often used for numerous cell biology experiments where cell delivery is involved (delivery of fluorescent probes, transfection of DNA, gene editing, etc). Our goal was therefore to identify parameters that may improve outcomes (or generate variability) in these experiments as well.

Our initial experiments established that the flow cytometry analysis of cells incubated with TMR-TAT is a robust way to quantify uptake. Comparison between technical replicates yielded a low chi-square value (<40). Conversely, the dose-response established with TMR-TAT concentrations suggested that changes in uptake could be detected with a relatively good resolution. Biological triplicates (experiments carried out using the same protocol but performed on different days) were much less reproducible (data not shown, chi-square values >200). It should be noted that our assay includes a 1 hour incubation of cells with TMR-TAT. We therefore measure the continuous accumulation of the CPP within endosomes during this time period, as this would be generally useful for a delivery experiment. This is in contrast with studies that measure the uptake of cell surface-bound ligands while washing away excess non-bound ligand (typically using 4°C to block uptake during ligand binding).[47] It is possible that the 1 h incubation, and the continuous uptake of CPPs during this time frame obscures some fluctuations that may happen on faster time scales. One should also note that the concentration of CPP typically used (5 μM) represents a large excess in comparison to cells (∼10 billion peptides per cell for a well containing 60,000 cells and 200 μL incubation media). It has been reported that the polyarginine CPP R12 has the capability to bind the C-X-C chemokine receptor type 4 (CXCR4) on the cell surface and that this binding stimulates macropinocytic uptake.[48] It is possible that TAT, in addition to the binding to heparin sulfate proteoglycans described in introduction, also shows some binding preference towards specific receptors. However, given that the abundance of a specific transmembrane receptor is usually rather low (10^2^-10^6^/cell), it is likely that the stoichiometry of the assay favors the binding of TMR-TAT to many different cell surface components.[49, 50] The uptake detected in our experiments are likely the result of TAT-cell membrane interactions and overall changes in macropinocytosis rate, which as previously described predominate over minor contributors to endocytosis, such as receptor mediated endocytosis. Hence these results do not reflect subtle variations that may occur. By surveying several intrinsic and extrinsic factors that may modulate endocytosis, we have identified parameters that do not perturb TMR-TAT uptake and parameters that modulate this activity. Cell density, potential stress caused due to washing steps, and cell aging, did not have a detectable impact on TMR-TAT. This is despite previous reports suggesting that cell-to-cell contact reduces endocytic uptake, that shear stress enhances it, and that it is downregulated in senescent cells [51-54]. The previous reports highlighted effects on receptor-mediated endocytosis. Because the bulk of TMR-TAT is likely endocytosed primarily by macropinocytosis, it is possible that these factors do not impact this uptake pathway. In contrast, the dissociation reagent, the composition of growth and incubation media, serum starvation, and temperature were factors that all had a detectable impact on uptake. The simple step of trypsinization, a routine protocol in cell culturing of adherent cells, was a major contributor to this variability, to a greater extent than the CDB approach. This is consistent with the notion that trypsin cleaves and removes numerous cell surface proteins, hence leading to a pronounced reorganization of the plasma membrane [26]. The recovery processes involved in repairing this damage may be somewhat chaotic, leading to different cell surface compositions from one cell culture to another, and hence yield variable endocytosis responses.

The formulation of the incubation media, and the concentration of glucose in particular, also had an impact on uptake. Glucose being intimately linked to metabolism, it is likely that its effects on endocytosis are pleiotropic. Notably, high glucose concentrations stimulate the endocytosis of glucose transporters, a phenomenon that regulates glucose uptake (by removing glucose transporters from its plasma membrane, cells lower glucose access to their cytoplasm) [54]. It is possible that TMR-TAT uptake is piggy backing this process. The effect of growth media was either not detected or small, possibly indicating that cells adapt rapidly to media changes (at least within the 1h incubation time frame). It should be noted that, during incubation, it is possible that it is not the rate of macropinosome formation and internalization that differ between media conditions. Instead, components that differ between media could potentially interfere with binding of TMR-TAT to the cell surface, thereby diminishing its uptake. Overall, it is nonetheless clear that the choice of media used for CPP delivery protocols is a critical consideration for experimental design.

Endocytic rates are temperature dependent [47, 55-57]. Endocytic uptake is generally negligible at temperatures below 20°C, and increasing steadily between 20 and 40°C as demonstrated on different cell types with different probes [47, 55-58]. To our knowledge, while blocking uptake at 4°C is often used as a means to test mechanisms of CPP penetration, the general effect of temperature on CPP uptake have not been explored. On one hand, it is clear that there is no obvious reason to carry out a cell incubation at 25°C, when it can easily be done in an incubator at 37°C. On the other, keeping practical considerations in mind, it is in actuality challenging to keep cells at a constant temperature of 37°C during cell culturing steps. For instance, prior to incubation, cells are handled in a biosafety laminar flow hood at room temperature, and washed and exposed to media that may or may not be at 37°C. It is also unclear how long it takes for the temperature to equilibrate when cell dishes are moved from incubator to biosafety cabinet and vice versa. Instead of testing the effect of transient temperature changes, we chose to test TMR-TAT uptake in cells maintained at 25°C throughout an experiment. We expected that this condition would represent a potential “worst-case” scenario that would help establish a lower boundary for the effect of room temperature on uptake. The rate of TMR-TAT uptake at 25°C was found to be approximately 50% of that obtained at 37°C. In turn, this indicates that cells exposed to room temperature for variable lengths of time may yield significant differences in uptake. For instance, using the equations provided in Figure 4, one can anticipate that cells exposed to 25°C for 5 or 10 min would endocytose 5% or 9% less peptide respectively than cells maintained at 37°C throughout a 1 hour incubation. In practice, it may be possible to increase the incubation temperature to 40°C to further increase uptake. However, given that heat shock stress can elicit numerous deleterious responses in cells, we did not pursue testing this condition [59].

While the factors described above contribute to small to moderate changes in CPP uptake individually, it is their combination that yielded the largest impact. As described above, the LUP tested was not necessarily representative of a procedure that would be typically used in a delivery experiment. Instead, along with the MUP and LUP, it was used to establish some practical boundaries that reflect the overall range by which endocytic uptake may vary between a worst and best-case scenario. The results obtained from the three protocols show significant differences in overall uptake, specifically in the number of TMR-TAT containing puncta per cell (indicative of the number of endosomes containing TMR-TAT). Together, these results indicate that the amount of a CPP that accumulates within endocytic organelles can vary in concentration by an order of magnitude depending on the protocol used. They also show that the different parameters used in a protocol have a cumulative effect (e.g. the difference seen between MUP and HUP, 2.3-fold MFI change, is higher than the 1.4-fold change observed between for the conditions comparing L-15 and DMEM). Given that there is a broad range of bulk endocytosis detected using our different protocols and that delivery efficiency is related to the amount of endocytosed CPP, we suspect that this in part may explain why CPPs may exhibit variable delivery efficiencies depending on the reported study, as protocols may be slightly different depending on the application. In particular, the microscopy results suggest that depending on the protocol used, TMR-TAT enters more endocytic compartments, rather than increasing number of TMR-TAT molecules per compartment. It is possible that different protocols could stimulate the formation of macropinosomes, resulting in an increased number of endosomes being present in our HUP, allowing for more TMR-TAT to enter the endocytic pathway. Hence, it is reasonable to speculate that the differences between HUP and MUP (resulting from serum starvation and differences in media components) could be due to alterations in metabolic states that result in changes in how many endosomes the cells produce and maintain at homeostatis. Alternatively, the rate by which TMR-TAT distributes throughout endosomal organelles may be more rapid and more efficient for the HUP conditions. Elucidating these various scenarios will require further studies. At this point, it is nonetheless clear that an increase in uptake is not synonymous with increased luminal concentration within a given endosome. One could therefore envision that TMR-TAT is not more likely to escape a endosome from cells in the HUP over the MUP because the CPP luminal concentrations are equivalent. Instead, one can also envision that the more TMR-TAT loaded endosomes a cell contains, the more likely it is that one or more of these endosomes will leak. Overall, the results presented highlight the importance of identifying protocol steps that impact endocytosis, either positively or negatively. This can be useful to improve experimental reproducibility. This can also be exploited to potentially increase uptake and, as highlighted herein, the number of endosomes that contain delivery agents. In future studies, we will test whether improving endocytic uptake will also directly or indirectly improve endosomal escape, and hence overall cell penetration and payload delivery. Finally, while several of the cell culturing parameters identified herein are not relatable to *in vivo* settings, it is intriguing to speculate that some factors, glucose concentration for instance, may be exploitable in some translational applications.

## Materials and Methods

### Cell Culture

HeLa (ATCC CCL-2), was cultured in Dulbecco’s minimum essential media (DMEM) (Fisher) supplemented with 10% fetal bovine serum (FBS) (Fisher), unless specifically noted for particular experiments. Cells were grown in a 37°C incubator with 5% CO_2_.

### Synthesis and purification of TMR-TAT

TMR-TAT was synthesized and purified as previously described (3). Briefly, 150 mg of rink-amide MBHA (0.5 mmol/g loading) was swelled in DMF for 30 minutes. The Fmoc group on the rink-amide MBHA (Bachem) resin was removed with 20% piperidine, yielding a free amine. This amine was reacted with an activated amino acid using the following cocktail: fmoc-gly-OH (3.9 eq), HCTU (4.0 eq), DIEA (10 eq). This process was repeated with the appropriate amino acid until the TAT sequence^49-58^ (RKKQRRRG) was completed. 5(6)-Carboxytetramethylrhodamine was then coupled to the N-terminus using the following cocktail: 5(6)-Carboxytetramethylrhodamine (2 eq), HCTU (2 eq), DIEA (5 eq). TMR-TAT was then cleaved from the resin using a mixture of 95%TFA/2.5% water/2.5%TIS. TMR-TAT concentration was determined by measuring the absorbance at 556 nm (91,500 cm^-1^ M^-1^) and was further confirmed by amino acid analysis (AAA). The N-terminally modified arginine (amino acid position 49) was excluded from AAA. To analyze the exact mass of the peptide, 10 µL of a 5 µM sample was adsorbed onto an α-cyano-4-hydroxycinnamic acid (CHCA) matrix (sigma) and subsequently analyzed via a Bruker Ultraflex Xtreme Maldi-TOF.

### Measurements for endocytic uptake by flow cytometry

Dried TMR-TAT powder was first weighed, and the weight was used to estimate the amount of water needed to resuspend the peptide solution to a concentrated solution (approximately 1 mM). This solution was then measured for concentration and was diluted to a make separate TMR-TAT stocks of either 40, 80, 120, 160, 200, 400 µM of TMR-TAT. These TMR-TAT solutions were diluted to the appropriate concentration (5 µL for 1-10 µM in 200 µL L-15) depending on the assay. For most assays 5 µM of TMR-TAT was utilized unless specifically stated otherwise. In the case of actin polymerization inhibition using cytochalasin D (cytoD), cells were preincubated for 30 minutes with 20 µM of cytoD. TMR-TAT was then added to cells in the presence 20 µM of CytoD. To prepare the cells for TMR-TAT treatment, cells were washed 3 times with Leibovitz’s L-15 to remove DMEM supplemented with 10% FBS, as components in FBS are suspected to interact with TAT-like molecules, hence disrupting endocytosis and/or delivery, and FBS may contain additional proteases that may degrade arginine rich peptides [31, 60]. The cells were then incubated at 37°C for 1 hour, unless otherwise specified (see figure legends). To wash away excess peptide, cells were then washed three times with L-15 supplemented with heparin (1 mg/mL) to remove extracellular peptide. To dissociate cells for flow cytometry, cells were removed by Trypsin (0.5% v/v in PBS) treatment, which cleaves extracellular proteins and likely removes any remaining TMR-TAT bound to the surface of cells (figure 2). Cells were then resuspended in L-15 medium and were kept on ice until analysis. Fluorescence signal of the TMR-TAT was detected on a flow cytometer (BD Accuri C6 model) using a standard FL2 (λex/λem = 585/640 ± 30 nm) channel. Data were acquired at a flow rate of 66 μL/min with at least 10,000 events. The fluorescence signals were processed in FlowJo v10.8. The initial experiment to establish this assay was performed in triplicate (see figure 1), while all other assays were performed in duplicate on the same day.

### Live cell imaging

Cells were imaged using an EVOSm7000 confocal microscope (20X objective) or an Olympus IX70 confocal microscope (100x images) with the appropriate filter cubes.

### SDS-PAGE and in gel fluorescence

From a 48 well dish, cells were solubilized using 100 µL of lysis buffer (1% TX-100, 1X-Halt protease inhibitor cocktail, 20 mM Tris, 150 mM NaCl, pH 7.4). 5X SDS-loading dye was immediately added to the sample. The sample was then boiled for 5 minutes in a 100°C dry-bath. Samples were then run on a commercially available 16% SDS-PAGE gels (Thermofisher) at 150 V for approximately 1 hour. NHS-ester and BSA Alexa Fluor™ 488 labeled conjugates (Thermofisher) were detected using a typhoon imager using a blue laser 473 nm and a LPB filter type (>510 nm).

### Colocalization of TMR-TAT with lysotracker

Cells were treated with TMR-TAT as described above. Subsequently, cells were treated with a cocktail containing lysotracker 0.5 µM and 1.5 µM of Hoeschst 33342 trihydrochloride trihydrate dye in L-15 for 15 minutes. This experiment was duplicated, and per replicate, 40-50 cells were quantified for colocalization of TMR-TAT and lysotracker using an imageJ plug-in called JaCoP [61].

### Comparison of cell detachment methods from cells

Cells were dissociated for 5 minutes with either trypsin (0.5% v/v) or an enzyme-free cell dissociation buffer (CDB) (Thermofisher). Although the exact composition is not disclosed by Thermofisher, it is known that CDB contains many chelators as the primary dissociation agent. Chelators allow for the dissociation of cells as many membrane proteins responsible for adherence require metal cofactors [27]. Cells were then allowed to proceed growth for 4 hours (minimum amount of time required for a majority of the cells to become adhered), and 24 hours (typical recovery time before performing an experiment). At these time points, cells were treated with TMR-TAT as previously described. In parallel, cells were labeled with Alexa-Fluor 488™ NHS-ester to detect cell surface proteins at each time point. To perform the Alexa-Fluor 488TM NHS-ester labeling, cells were washed three times with Hanks balanced salt solution (HBSS buffer). A 1.9 mM stock solution (dry powder dissolved in DMSO) of Alexa FluorTM 488 NHS-ester (Thermo-fisher) was diluted to 1.9 µM in HBSS buffer. The Alexa Fluor 488™ NHS-ester (1.9 µM) in HBSS was added directly to live cells. The amine labeling of cell proteins was allowed to proceed for 30 minutes at 37°C. Cells were then washed 3 times with HBSS buffer to remove any detached cells. The adherent cells were lysed with lysis buffer and run on SDS-PAGE analysis as previously described.

### Treatment of cells with Alexa Fluor488™ BSA conjugate

Alexa Fluor 488™ BSA conjugate was used to assess the amount of BSA remaining after subsequent washes. To execute the experiment, approximately 50,000 cells were plated on a 48 well plate and cultured for 24 hours under standard conditions, as described above. Alexa Fluor 488™ BSA conjugate (5 mg) was resuspended in PBS to a concentration of 150 µL. This stock solution was then diluted in DMEM supplemented with 10% fetal bovine serum to a concentration of 3.25 µM. 24 hours later, cells were then washed a variable amount of times, and then lysed (as previously described). The cell lysates were then run on SDS-PAGe and analyzed for fluorescence via a typhoon laser scanner, as previously described.

### Comparison of media on TMR-TAT uptake

Cells were dissociated with 0.5% trypsin. Trypsin was then inactivated by either high glucose DMEM (HG-DMEM) pyruvate with 10% FBS or low glucose DMEM (LG-DMEM) with 10% FBS. HG-DMEM (25 mM glucose) was purchased from Thermofisher. LG-DMEM was prepared by adding Glucose (5 mM) to non-glucose containing DMEM. Cells were then seeded onto a 48 well with 50,000 cells per well and were allowed to grow for exactly 24 hours. Cells were then prepared for TMR-TAT treatment (5 µM), as previously described, except TMR-TAT was diluted into either Leibovitz’s L-15 (used for previously described experiments unless specifically noted) without 10% FBS, DMEM-HG without 10% FBS, or DMEM-LG without 10% FBS. After 1 hour incubation of TMR-TAT, cells were prepared for flow cytometry, as previously described.

### Measuring the impact of media exchange and serum starvation

Cells were washed three times with the selected buffer where the cells would undergo serum starvation (either Leibovitz’s L-15 without 10% FBS or DMEM-HG). In separate wells, cells were then serum starved for 0, 1.5, and 3.0 hours. Subsequently, cells were treated with TMR-TAT (5 µM) for 1 hour and were then collected for flow cytometry analysis.

### Temperature dependence assay for TMR-TAT uptake

Cells were plated into two separate 48 well plates. For each well plate, six wells were seeded, one for each timepoint (5 min, 15 min, 30 min, 45 min, 60 min, untreated). After 24 hours of growth, each well plate was equilibrated for 15 minutes to the desired temperature (25°C or 37°C). Cells were then prepared for TMR-TAT incubation. Throughout the TMR-TAT incubation, cells were kept on either a 25°C or on a 37°C heat stage. At each time point, cells were washed with 1X heparin, dissociated with trypsin, and analyzed by flow cytometry as previously described.

### Comparison of low uptake (LUP), medium uptake (MUP) and high uptake protocol (HUP)

For all protocols, cell were dissociated with CDB, since it is least likely to yield variable results as compared to trypsin (see figure 2). After 24 hours of growth, cells grown in HG-DMEM containing 10% FBS (LUP and MUP) or LG-DMEM (HUP) was removed and washed three times with either Leibovitz’s L-15 without FBS (LUP and MUP) or DMEM without FBS (HUP). For both LUP and MUP, cells were then incubated for 3 hours in Leibovitz’s L-15 without FBS. For LUP, cells were then equilibrated to 25°C for 15 minutes and then treated with TMR-TAT (stock solution diluted in Leibovitz’s L-15 without FBS to make final concentration of 5 µM) and incubated for one hour. For MUP, the same TMR-TAT treatment procedures were conducted, but at 37°C instead. For the high uptake protocol, cells were treated with TMR-TAT (stock solution diluted in HG-DMEM without FBS to make final concentration of 5 µM) and incubated for one hour at 37°C. Cells were then collected for flow cytometry analysis. In parallel, cells were visualized by 100x microscopy. Using Slidebook 6 to control the microscope, it was ensured that all images were taken on the same focal plane. Slidebook 6 was also used to deconvolute images to eliminate out of focus endosomes on different planes. In Celleste imaging analysis software, the average pixel intensities of individual objects containing TMR-TAT was collected and used as a proxy to compare potential differences in the amount of TMR-TAT in endosomal compartments. Additionally, single cells (total 50 each per condition) were selected using the Celleste software region of interest (ROI) selection tool, and the number of pixels per cell was collected. The number of pixels per cell is proportional to the number of TMR-TAT containing endosomes. It is important to note that this measurement does not give a direct number of endosomes per cell, since individual endosomes may not be fully resolved by microscopy and only one plane of the cell is assessed. Despite these limitations, these data can be used to make comparisons of how the relative number of TMR-TAT containing puncta changes among LUP, MUP, and HUP. To illustrate these measurements, surface plots corresponding to the pixel intensities of each puncta containing TMR-TAT was generated in ImageJ plugin interactive 3D surface plot.

## Supplemental methods

### Impact of cell density on endocytosis

The density of cells adhering to a surface, or whether cells make contact with one another, has been reported to impact endocytic uptake [51]. In order to test how potential cell stress impacts the uptake of TMR-TAT, cells were seeded at different dilutions during passaging and allowed to grow overnight. To quantify cell density, cells were stained with Hoechst (1.25 µM) and imaged by fluorescence microscopy (over the whole area of the dish to ensure homogeneity of distribution). Confluent cells were used to establish a maximal density of blue nuclei. This was then used to determine a ratio of surface density (Figure S2). Cells were also stained with di-4-ANEPPDHQ (5.0 µM), a lipophilic probe, to label their membranes and to confirm that cells contact each other at high density [30]. Cells at various densities were then incubated with TMR-TAT using SP and analyzed by flow cytometry. The populations of TMR-TAT positive cells were indistinguishable in all conditions tested, as indicated by Chi squared T(x) scores of zero (Figure S2).

### Cell washing protocol

Cell washing is required to remove culture medium used for cell growth and to introduce of a medium appropriate for a delivery experiment. In the case study described herein, this involves removing phenol red-DMEM, which is a medium using a sodium bicarbonate buffering system adapted to growth in 5% CO_2_ incubators, and replacing it with Leibovitz’s L-15, a medium buffered by phosphates and compatible with ambient air (and hence better suited for microscopy experiments). Despite being relatively trivial, media exchange can be stressful to cells. In particular, flow and shear stress can impact cell physiology, both in vivo and in vitro [34-37]. To test how cell washing impacts TMR-TAT endocytosis, cells were washed by following three distinct protocols: dry, aspirate, dilution. The dry technique involves removing media with a pipette while aspirate uses an aspirator connected to a vacuum. In both techniques, most of the visible liquid is removed, leaving liquid at the corner of the dish (∼ 20 µL). The dilution technique is similar to the dry technique with the exception that a layer of liquid (200 µL) is left in a dish prior to introduction of volume of new media. In all cases, new media is added slowly with a pipette on the side corner of a well, so as to avoid additional stress. The density of cells attached to a dish, along with their viability, were measured before and after each cell washing protocol. The aspirate and dilution protocols increased the number of cells that detached from the dish when compared to the dry protocol (it should be noted that the aspirate protocol can have a detrimental effect on the number of cells that remain attached to cells if the aspirator is left in contact with cells for an extended period of cells, ∼>5 sec; not shown). Conversely, the dry and aspirate protocols led to the presence of more dead cells remaining in the dish than the dilution protocol. Nonetheless, despite evidence that cells experience different levels of stress, the uptake of TMR-TAT was not impacted by these parameters, as indicated by low Chi squared T(x) scores in flow cytometry results (Figure S3).

### Washing steps

Albumin present in the supplement FBS has been shown to be inhibitory to TAT cell penetration. This is likely because albumin, a negatively charged protein at pH 7 (pI = 4.9), binds the cationic peptide and prevents it from interacting with other cell components. Albumin can bind the cell surface and adsorb to labware (BSA can for instance be used to coat plastic or glass and prevent adsorption of other proteins) [62]. To assess how the presence of albumin (or other components present in FBS) may impact TMR-TAT endocytosis, the incubation media used for cell growth (10% FBS supplemented DMEM) was exchanged to TMR-TAT/L15 media, directly (0 wash) or after 1, 2, and 3 washes with L15. The amount of albumin left in the dish after each wash was assessed by SDS-PAGE. Alternatively, to detect low levels of albumin, FITC-albumin was used instead and detected by SDS-PAGE and fluorescence imaging. Flow cytometry analysis of TMR-TAT endocytosis under these conditions show that direct exchange of media leads to a population of cells with lower endocytic uptake (as indicated by a median lower than the mode). Cells washed 1 to 3 times and treated with TMR-TAT were otherwise indistinguishable (Figure S4).

### Impact of cell culture age of TMR-TAT uptake

Cells were passaged up to 25 days. On day 8 and day 23 of passaging, cells were dissociated with CDB and spun at 500 x g to collect the cell pellet. The supernatant was removed, and the cell pellet was resuspended in DMEM/10% DMSO (supplemented with 10% FBS). Cell stocks were then stored in a liquid nitrogen dewar (–196°C). On the same day, the original stock (day 0), and stocks stored on day 8 and day 23 were revived. Cells stocks were passaged twice and seeded in 48 well dishes. Cells were then treated with TMR-TAT (5 µM) for one hour and subsequently analyzed by flow cytometry.

## Statements and Declarations

### Author Contributions

All authors contributed to the design of the experiments and preparation of the manuscript.

### Data Availability

As applicable, data sets will be made available if needed upon review for publication.

### Ethics

Not applicable

### Consent to participate

Not applicable

### Consent to publish

Not applicable

DBmz8f$KM&tkaS8z

## References

1. G. J. Doherty and H. T. McMahon Mechanisms of Endocytosis Annual Review of Biochemistry 78 (2009) 857–902. DOI: 10.1146/annurev.biochem.78.081307.110540.

2. A. K. Varkouhi, M. Scholte, G. Storm and H. J. Haisma Endosomal escape pathways for delivery of biologicals Journal of Controlled Release 151 (2011) 220–228. DOI: https://doi.org/10.1016/j.jconrel.2010.11.004.

3. D. Pei and M. Buyanova Overcoming Endosomal Entrapment in Drug Delivery Bioconjugate chemistry 30 (2019) 273–283. DOI: 10.1021/acs.bioconjchem.8b00778.

4. S. L. Y. Teo, J. J. Rennick, D. Yuen, H. Al-Wassiti, A. P. R. Johnston and C. W. Pouton Unravelling cytosolic delivery of cell penetrating peptides with a quantitative endosomal escape assay Nature Communications 12 (2021) 3721. DOI: 10.1038/s41467-021-23997-x.

5. A. Bolhassani, B. S. Jafarzade and G. Mardani In vitro and in vivo delivery of therapeutic proteins using cell penetrating peptides Peptides 87 (2017) 50–63. DOI: https://doi.org/10.1016/j.peptides.2016.11.011.

6. M. Silhol, M. Tyagi, M. Giacca, B. Lebleu and E. Vivès Different mechanisms for cellular internalization of the HIV-1 Tat-derived cell penetrating peptide and recombinant proteins fused to Tat Eur J Biochem 269 (2002) 494–501. DOI: 10.1046/j.0014-2956.2001.02671.x.

7. M. Tyagi, M. Rusnati, M. Presta and M. Giacca Internalization of HIV-1 tat requires cell surface heparan sulfate proteoglycans J Biol Chem 276 (2001) 3254–3261. DOI: 10.1074/jbc.M006701200.

8. J. P. Richard, K. Melikov, H. Brooks, P. Prevot, B. Lebleu and L. V. Chernomordik Cellular uptake of unconjugated TAT peptide involves clathrin-dependent endocytosis and heparan sulfate receptors J Biol Chem 280 (2005) 15300–15306. DOI: 10.1074/jbc.M401604200.

9. E. Gonçalves, E. Kitas and J. Seelig Binding of Oligoarginine to Membrane Lipids and Heparan Sulfate: Structural and Thermodynamic Characterization of a Cell-Penetrating Peptide Biochemistry 44 (2005) 2692–2702. DOI: 10.1021/bi048046i.

10. I. Nakase, K. Osaki, G. Tanaka, A. Utani and S. Futaki Molecular interplays involved in the cellular uptake of octaarginine on cell surfaces and the importance of syndecan-4 cytoplasmic V domain for the activation of protein kinase Calpha Biochem Biophys Res Commun 446 (2014) 857–862. DOI: 10.1016/j.bbrc.2014.03.018.

11. J. F. Casella, M. D. Flanagan and S. Lin Cytochalasin D inhibits actin polymerization and induces depolymerization of actin filaments formed during platelet shape change Nature 293 (1981) 302–305. DOI: 10.1038/293302a0.

12. M. Koivusalo, C. Welch, H. Hayashi, C. C. Scott, M. Kim, T. Alexander, N. Touret, K. M. Hahn and S. Grinstein Amiloride inhibits macropinocytosis by lowering submembranous pH and preventing Rac1 and Cdc42 signaling J Cell Biol 188 (2010) 547–563. DOI: 10.1083/jcb.200908086.

13. H. Hackstein, T. Taner, A. J. Logar and A. W. Thomson Rapamycin inhibits macropinocytosis and mannose receptor-mediated endocytosis by bone marrow-derived dendritic cells Blood 100 (2002) 1084–1087. DOI: 10.1182/blood.v100.3.1084.

14. F. Madani, S. Lindberg, U. Langel, S. Futaki and A. Gräslund Mechanisms of cellular uptake of cell-penetrating peptides J Biophys 2011 (2011) 414729. DOI: 10.1155/2011/414729.

15. J. P. Lim and P. A. Gleeson Macropinocytosis: an endocytic pathway for internalising large gulps Immunol Cell Biol 89 (2011) 836–843. DOI: 10.1038/icb.2011.20.

16. J. Klumperman and G. Raposo The complex ultrastructure of the endolysosomal system Cold Spring Harbor perspectives in biology 6 (2014) a016857–a016857. DOI: 10.1101/cshperspect.a016857.

17. I. Mäger, K. Langel, T. Lehto, E. Eiríksdóttir and Ü. Langel The role of endocytosis on the uptake kinetics of luciferin-conjugated cell-penetrating peptides Biochimica et Biophysica Acta (BBA) - Biomembranes 1818 (2012) 502–511. DOI: https://doi.org/10.1016/j.bbamem.2011.11.020.

18. J. K. Allen, D. J. Brock, H. M. Kondow-McConaghy and J. P. Pellois Efficient Delivery of Macromolecules into Human Cells by Improving the Endosomal Escape Activity of Cell-Penetrating Peptides: Lessons Learned from dfTAT and its Analogs Biomolecules 8 (2018). DOI: 10.3390/biom8030050.

19. A. Mishra, H. Lai Ghee, W. Schmidt Nathan, Z. Sun Victor, R. Rodriguez April, R. Tong, L. Tang, J. Cheng, J. Deming Timothy, T. Kamei Daniel and C. L. Wong Gerard Translocation of HIV TAT peptide and analogues induced by multiplexed membrane and cytoskeletal interactions Proceedings of the National Academy of Sciences 108 (2011) 16883–16888. DOI: 10.1073/pnas.1108795108.

20. S. G. Patel, E. J. Sayers, L. He, R. Narayan, T. L. Williams, E. M. Mills, R. K. Allemann, L. Y. P. Luk, A. T. Jones and Y.-H. Tsai Cell-penetrating peptide sequence and modification dependent uptake and subcellular distribution of green florescent protein in different cell lines Scientific Reports 9 (2019) 6298. DOI: 10.1038/s41598-019-42456-8.

21. D. Mittelman and J. H. Wilson The fractured genome of HeLa cells Genome Biology 14 (2013) 111. DOI: 10.1186/gb-2013-14-4-111.

22. M. Roederer and R. R. Hardy Frequency difference gating: a multivariate method for identifying subsets that differ between samples Cytometry 45 (2001) 56–64. DOI: 10.1002/1097-0320(20010901)45:1<56::aid-cyto1144>3.0.co;2-9.

23. M. Roederer, W. Moore, A. Treister, R. R. Hardy and L. A. Herzenberg Probability binning comparison: a metric for quantitating multivariate distribution differences Cytometry 45 (2001) 47–55. DOI: 10.1002/1097-0320(20010901)45:1<47::aid-cyto1143>3.0.co;2-a.

24. M. Roederer, A. Treister, W. Moore and L. A. Herzenberg Probability binning comparison: a metric for quantitating univariate distribution differences Cytometry 45 (2001) 37–46. DOI: 10.1002/1097-0320(20010901)45:1<37::aid-cyto1142>3.0.co;2-e.

25. M. C. Phelan Basic Techniques in Mammalian Cell Tissue Culture Current Protocols in Cell Biology 36 (2007) 1.1.1-1.1.18. DOI: https://doi.org/10.1002/0471143030.cb0101s36.

26. H. L. Huang, H. W. Hsing, T. C. Lai, Y. W. Chen, T. R. Lee, H. T. Chan, P. C. Lyu, C. L. Wu, Y. C. Lu, S. T. Lin, C. W. Lin, C. H. Lai, H. T. Chang, H. C. Chou and H. L. Chan Trypsin-induced proteome alteration during cell subculture in mammalian cells J Biomed Sci 17 (2010) 36. DOI: 10.1186/1423-0127-17-36.

27. B. Leitinger, A. McDowall, P. Stanley and N. Hogg The regulation of integrin function by Ca2+ Biochimica et Biophysica Acta (BBA) - Molecular Cell Research 1498 (2000) 91–98. DOI: https://doi.org/10.1016/S0167-4889(00)00086-0.

28. E. J. O’Keefe, R. A. Briggaman and B. Herman Calcium-induced assembly of adherens junctions in keratinocytes J Cell Biol 105 (1987) 807–817. DOI: 10.1083/jcb.105.2.807.

29. W. Meng and M. Takeichi Adherens junction: molecular architecture and regulation Cold Spring Harbor perspectives in biology 1 (2009) a002899–a002899. DOI: 10.1101/cshperspect.a002899.

30. X. Zhao, R. Li, C. Lu, F. Baluška and Y. Wan Di-4-ANEPPDHQ, a fluorescent probe for the visualisation of membrane microdomains in living Arabidopsis thaliana cells Plant Physiology and Biochemistry 87 (2015) 53–60. DOI: https://doi.org/10.1016/j.plaphy.2014.12.015.

31. J. Allen, K. Najjar, A. Erazo-Oliveras, H. M. Kondow-McConaghy, D. J. Brock, K. Graham, E. C. Hager, A. L. J. Marschall, S. Dubel, R. L. Juliano and J. P. Pellois Cytosolic Delivery of Macromolecules in Live Human Cells Using the Combined Endosomal Escape Activities of a Small Molecule and Cell Penetrating Peptides ACS Chem Biol 14 (2019) 2641–2651. DOI: 10.1021/acschembio.9b00585.

32. W.-H. Tang, C.-F. Wang and Y.-D. Liao Fetal bovine serum albumin inhibits antimicrobial peptide activity and binds drug only in complex with α1-antitrypsin Scientific Reports 11 (2021) 1267. DOI: 10.1038/s41598-020-80540-6.

33. L. T. Nguyen, J. K. Chau, N. A. Perry, L. de Boer, S. A. Zaat and H. J. Vogel Serum stabilities of short tryptophan- and arginine-rich antimicrobial peptide analogs PLoS One 5 (2010). DOI: 10.1371/journal.pone.0012684.

34. B. Sinha, D. Köster, R. Ruez, P. Gonnord, M. Bastiani, D. Abankwa, R. V. Stan, G. Butler-Browne, B. Vedie, L. Johannes, N. Morone, R. G. Parton, G. Raposo, P. Sens, C. Lamaze and P. Nassoy Cells Respond to Mechanical Stress by Rapid Disassembly of Caveolae Cell 144 (2011) 402–413. DOI: 10.1016/j.cell.2010.12.031.

35. F. Boccafoschi, M. Bosetti, P. M. Sandra, M. Leigheb and M. Cannas Effects of mechanical stress on cell adhesion: a possible mechanism for morphological changes Cell Adh Migr 4 (2010) 19–25. DOI: 10.4161/cam.4.1.9569.

36. Y.-M. Lin, F. Li and X.-Z. Shi Mechanical Stress Is a Pro-Inflammatory Stimulus in the Gut: In Vitro, In Vivo and Ex Vivo Evidence PLOS ONE 9 (2014) e106242. DOI: 10.1371/journal.pone.0106242.

37. G. Nagamatsu, S. Shimamoto, N. Hamazaki, Y. Nishimura and K. Hayashi Mechanical stress accompanied with nuclear rotation is involved in the dormant state of mouse oocytes Sci Adv 5 (2019) eaav9960. DOI: 10.1126/sciadv.aav9960.

38. R. Shiratori, K. Furuichi, M. Yamaguchi, N. Miyazaki, H. Aoki, H. Chibana, K. Ito and S. Aoki Glycolytic suppression dramatically changes the intracellular metabolic profile of multiple cancer cell lines in a mitochondrial metabolism-dependent manner Sci Rep 9 (2019) 18699. DOI: 10.1038/s41598-019-55296-3.

39. C. N. Antonescu, T. E. McGraw and A. Klip Reciprocal regulation of endocytosis and metabolism Cold Spring Harb Perspect Biol 6 (2014) a016964. DOI: 10.1101/cshperspect.a016964.

40. S. Pirkmajer and A. V. Chibalin Serum starvation: caveat emptor Am J Physiol Cell Physiol 301 (2011) C272–279. DOI: 10.1152/ajpcell.00091.2011.

41. M.-u. Rashid and K. M. Coombs Serum-reduced media impacts on cell viability and protein expression in human lung epithelial cells Journal of Cellular Physiology 234 (2019) 7718–7724. DOI: https://doi.org/10.1002/jcp.27890.

42. R. Khammanit, S. Chantakru, Y. Kitiyanant and J. Saikhun Effect of serum starvation and chemical inhibitors on cell cycle synchronization of canine dermal fibroblasts Theriogenology 70 (2008) 27–34. DOI: https://doi.org/10.1016/j.theriogenology.2008.02.015.

43. E. Z. White, N. M. Pennant, J. R. Carter, O. Hawsawi, V. Odero-Marah and C. V. Hinton Serum deprivation initiates adaptation and survival to oxidative stress in prostate cancer cells Scientific Reports 10 (2020) 12505. DOI: 10.1038/s41598-020-68668-x.

44. A. Erazo-Oliveras, K. Najjar, L. Dayani, T. Y. Wang, G. A. Johnson and J. P. Pellois Protein delivery into live cells by incubation with an endosomolytic agent Nat Methods 11 (2014) 861–867. DOI: 10.1038/nmeth.2998.

45. A. J. M. Santos and E. Boucrot Probing Endocytosis During the Cell Cycle with Minimal Experimental Perturbation Methods Mol Biol 1847 (2018) 23–35. DOI: 10.1007/978-1-4939-8719-1_3.

46. C. Rosin, K. Estel, J. Hälker and R. Winter Combined Effects of Temperature, Pressure, and Co-Solvents on the Polymerization Kinetics of Actin ChemPhysChem 16 (2015) 1379–1385. DOI: https://doi.org/10.1002/cphc.201500083.

47. P. H. Weigel and J. A. Oka Temperature dependence of endocytosis mediated by the asialoglycoprotein receptor in isolated rat hepatocytes. Evidence for two potentially rate-limiting steps J Biol Chem 256 (1981) 2615–2617.

48. G. Tanaka, I. Nakase, Y. Fukuda, R. Masuda, S. Oishi, K. Shimura, Y. Kawaguchi, T. Takatani-Nakase, U. Langel, A. Graslund, K. Okawa, M. Matsuoka, N. Fujii, Y. Hatanaka and S. Futaki CXCR4 stimulates macropinocytosis: implications for cellular uptake of arginine-rich cell-penetrating peptides and HIV Chem Biol 19 (2012) 1437–1446. DOI: 10.1016/j.chembiol.2012.09.011.

49. D. Bausch-Fluck, A. Hofmann, T. Bock, A. P. Frei, F. Cerciello, A. Jacobs, H. Moest, U. Omasits, R. L. Gundry, C. Yoon, R. Schiess, A. Schmidt, P. Mirkowska, A. Hartlova, J. E. Van Eyk, J. P. Bourquin, R. Aebersold, K. R. Boheler, P. Zandstra and B. Wollscheid A mass spectrometric-derived cell surface protein atlas PLoS One 10 (2015) e0121314. DOI: 10.1371/journal.pone.0121314.

50. Y. Kawaguchi, T. Takeuchi, K. Kuwata, J. Chiba, Y. Hatanaka, I. Nakase and S. Futaki Syndecan-4 Is a Receptor for Clathrin-Mediated Endocytosis of Arginine-Rich Cell-Penetrating Peptides Bioconjug Chem 27 (2016) 1119–1130. DOI: 10.1021/acs.bioconjchem.6b00082.

51. B. Snijder, R. Sacher, P. Ramo, E. M. Damm, P. Liberali and L. Pelkmans Population context determines cell-to-cell variability in endocytosis and virus infection Nature 461 (2009) 520–523. DOI: 10.1038/nature08282.

52. E. Y. Shin, N. K. Soung, M. A. Schwartz and E. G. Kim Altered endocytosis in cellular senescence Ageing Res Rev 68 (2021) 101332. DOI: 10.1016/j.arr.2021.101332.

53. Z. He, W. Zhang, S. Mao, N. Li, H. Li and J. M. Lin Shear Stress-Enhanced Internalization of Cell Membrane Proteins Indicated by a Hairpin-Type DNA Probe Anal Chem 90 (2018) 5540–5545. DOI: 10.1021/acs.analchem.8b00755.

54. T. Lopez-Hernandez, V. Haucke and T. Maritzen Endocytosis in the adaptation to cellular stress Cell Stress 4 (2020) 230–247. DOI: 10.15698/cst2020.10.232.

55. N. L. Chanaday and E. T. Kavalali Time course and temperature dependence of synaptic vesicle endocytosis FEBS Lett 592 (2018) 3606–3614. DOI: 10.1002/1873-3468.13268.

56. I. Delvendahl, N. P. Vyleta, H. von Gersdorff and S. Hallermann Fast, Temperature-Sensitive and Clathrin-Independent Endocytosis at Central Synapses Neuron 90 (2016) 492–498. DOI: 10.1016/j.neuron.2016.03.013.

57. P. R. Sudhakaran, R. Prinz and K. Vonfigura Effect of Temperature on Endocytosis and Degradation of Sulfated Proteoglycans by Cultured Skin Fibroblasts J Bioscience 4 (1982) 413–418. DOI: Doi 10.1007/Bf02704633.

58. S. C. Silverstein, R. M. Steinman and Z. A. Cohn Endocytosis Annu Rev Biochem 46 (1977) 669–722. DOI: 10.1146/annurev.bi.46.070177.003321.

59. A. K. Velichko, E. N. Markova, N. V. Petrova, S. V. Razin and O. L. Kantidze Mechanisms of heat shock response in mammals Cell Mol Life Sci 70 (2013) 4229–4241. DOI: 10.1007/s00018-013-1348-7.

60. D. S. Youngblood, S. A. Hatlevig, J. N. Hassinger, P. L. Iversen and H. M. Moulton Stability of Cell-Penetrating Peptide−Morpholino Oligomer Conjugates in Human Serum and in Cells Bioconjugate Chemistry 18 (2007) 50–60. DOI: 10.1021/bc060138s.

61. S. Bolte and F. P. Cordelières A guided tour into subcellular colocalization analysis in light microscopy J Microsc 224 (2006) 213–232. DOI: 10.1111/j.1365-2818.2006.01706.x.

62. S.-M. Acuña-Nelson, J.-M. Bastías-Montes, F.-R. Cerda-Leal, J.-E. Parra-Flores, J.-S. Aguirre-García and P. G. Toledo Nanocoatings of Bovine Serum Albumin on Glass: Effects of pH and Temperature Journal of Nanomaterials 2020 (2020) 8640818. DOI: 10.1155/2020/8640818.

